# A simple, safe and sensitive method for SARS-CoV-2 inactivation and RNA extraction for RT-qPCR

**DOI:** 10.1101/2020.06.29.179176

**Authors:** Lelde Kalnina, Àngels Mateu-Regué, Stephanie Oerum, Annemette Hald, Jan Gerstoft, Henrik Oerum, Finn Cilius Nielsen, Astrid K.N. Iversen

## Abstract

The SARS-CoV-2 pandemic has created an urgent need for large amounts of diagnostic tests to detect viral RNA, which commercial suppliers are increasingly unable to deliver. In addition to the lack of availability, the current methods do not always fully inactivate the virus. Together, this calls for the development of safer methods for extraction and detection of viral RNA from patient samples that utilise readily available reagents and equipment present in most standard laboratories. We present a rapid and straightforward RNA extraction protocol for inactivating the SARS-CoV-2 virus that uses standard lab reagents. This protocol expands analysis capacity as the inactivated samples can be used in RT-qPCR detection tests at laboratories not otherwise classified for viral work. The method circumvents the need for commercial RNA purification kits, takes about 30 minutes from swab to PCR-ready viral RNA, and enables downstream detection of SARS-CoV-2 by RT-qPCR with very high sensitivity (~4 viral RNA copies per RT-qPCR). In summary, we present a rapid, safe and sensitive method for high-throughput detection of SARS-CoV-2, that can be conducted in any laboratory equipped with a qPCR machine.

## INTRODUCTION

In mid-December 2019, reports emerged that patients in Wuhan, Hubei province, China, were suffering from atypical pneumonia, and by start-January, the causative agent, severe acute respiratory syndrome coronavirus 2 (SARS-CoV-2), was identified (1, 2). The disease was named coronavirus disease 2019 (COVID-19). The initial epicentre of virus spread seems to have been the Huanan seafood wholesale market in Wuhan. Although the SARS-CoV-2 genome is very similar to bat SARS-CoV-like coronaviruses (~96%), it carries unique sequence motifs in the receptor-binding domain (RBD) of the Spike protein that binds to the human angiotensin-converting enzyme 2 (ACE2) receptor (3). These differences suggest that natural selection in an intermediate host species optimised binding of SARS-CoV-2 to ACE2, and facilitated transmission to, and spread between, humans. By mid-January 2020, the virus was found in Thailand and Japan following which it spread worldwide (4, 5). As of June 2020, the US had the largest number of identified SARS-CoV-2 infected individuals, but also several European countries, e.g., Italy, Spain, United Kingdom, and France have large numbers of COVID-19 patients (6). As of today, cases of COVID-19 are rapidly increasing in India, Mexico and parts of Africa and South America.

The possibility to rapidly test large numbers of individuals for the presence of SARS-CoV-2 is a vital component in containing viral spread, in understanding the infectious fatality rate, and in subsequently guiding the controlled reopening of our societies. In medical laboratories, the presence of SARS-CoV-2 is commonly detected in a two-step process, where each step requires different kits. Step one is the RNA extraction from patient swabs usually performed using a kit from Qiagen or Roche (7), and step two is the detection of SARS-CoV-2, often achieved by a reverse-transcriptase quantitative polymerase chain reaction (RT-qPCR). With the rapidly growing need for SARS-CoV-2 tests, commercial supplies are increasingly falling short on kits for both steps, thereby creating a need for alternative methods that utilise readily available reagents and equipment present in most standard laboratories.

A method, which combines a high molar acidic guanidinium isothiocyanate (GITC) solution, phenol and chloroform (collectively termed GPC), is broadly used to extract intact RNA from diverse biological samples (8, 9). The standard protocol for RNA-extraction by GCP is lengthy and requires significant expertise in handling RNA, making it unsuitable for large-scale screening programs. Here, we present a much-simplified version of the GPC-extraction method that overcomes these limitations while still providing inactivated, viral RNA that is compatible with downstream RT-qPCR detection. The method enables detection of ~4 viral RNA copies per RT-qPCR, corresponding to ~10^4^ viral RNA copies on the swab. This detection limit is substantially lower than the average virus load per nasopharyngeal (NP) or oropharyngeal (OP) swab from symptoms onset to day five (6.76×10^5^ copies per swab) and later during the disease course (3.44×10^5^) (10, 11). In addition to the protocol simplification, this modified method further offers safe working conditions for healthcare personnel as the GPC solution is known to rapidly and fully inactivate viruses of the corona family, e.g., MERS-CoV (12). The efficient viral inactivation enables the simplified GCP-extraction method to be used at laboratories not classified for viral work, when on-site capacity at authorised hospital laboratories presents an issue, thereby expanding testing capacity.

In summary, we propose a rapid, safe and sensitive method for high-throughput detection of SARS-CoV-2, that can be conducted in any laboratory equipped with an RT-qPCR machine, using inexpensive and readily available reagents.

## MATERIALS AND METHODS

### Protocol outline

The method comprises the following four steps.

1. Patient sampling by NP or OP swabs.
2. Addition of the GPC reagents that instantly inactivates the virus and protects the released viral RNA genome from enzymatic degradation.
3. Extraction of RNA in 2 simple steps.
4. RT-qPCR to detect SARS-CoV-2 RNA in the patient sample.

### Sample preparation and RNA extraction using the simplified GPC-extraction method

We obtained OP and NP samples from one hospitalised COVID-19 patient who had tested virus-positive three weeks earlier (cobas^®^ SARS-CoV-2 test, Roche diagnostics), from three individuals who had previously tested SARS-CoV-2 positive, but tested negative on the day of sampling (cobas^®^ SARS-CoV-2 test, Roche diagnostics), and from a healthy individual. NP and OP swabs were collected using FLOQSwabs^®^ (COPAN, 552C), placed in transport tubes, sealed and transferred to the laboratory. Each swab was incubated for 5 min at room temperature in an Eppendorf tube containing 1.1 mL TRI-reagent (T9424, Sigma), after which the swab was discarded, and 200 μL of chloroform was added. This sample/GPC mixture was vortexed for 30 sec, incubated for 3 min at room temperature, centrifuged for 15 min 12.000 g at 4°C, after which 10 μL of the upper aqueous phase was carefully transferred to a new tube without disturbing the interphase. The 10 μL sample was diluted with RNase-free water as detailed below.

### Dilution series of RNA extracted from OP and NP samples

To test the effect of GITC on the quality of both one-step and two-step RT-qPCR reactions, we created dilution series of the OP and NP samples that allowed the final dilution in the subsequent RT-reaction of the RT-qPCR to range from 8x to 100x (**Figure 1**). For example, to achieve a 100x final dilution in the RT-reaction of a 25 μL one-step RT-qPCR, the 10 μL sample was mixed with 190 μL RNase-free water to a dilution of 20x, followed by 5 μL of this sample mixed in a 25 μL one-step RT-qPCR to a final dilution of 100x.

**Figure 1.**
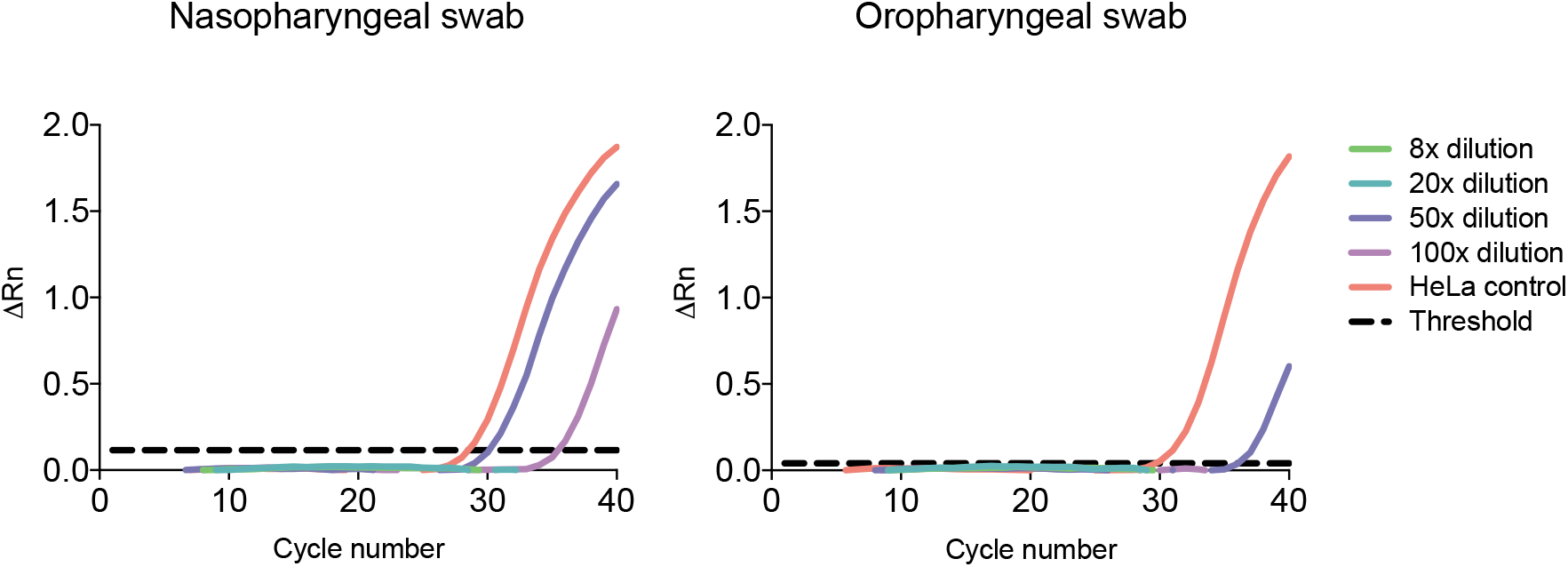
The effect of GITC salts on the efficiency of a two-step B2M RT-qPCR using dilutions of the aqueous phase from the GPC extraction of an NP and OP sample. A representative amplification plot of ΔRn against PCR cycle number for the two-step B2M RT-qPCR with different dilutions of the RNA-GITC solution in the RT reaction (ranging from 8x to 100x). No amplification was observed at 8x and 20x dilutions. The threshold is shown as a black dashed line and corresponds to 0.116 for the NP swab and 0.040 for the OP swab. ΔRn: Rn (the fluorescence of the reporter dye divided by the fluorescence of the passive reference dye ROX) minus the baseline (black dashed line). The amplifications were performed in duplicate.

### Two-step RT-qPCR against human B2M mRNA

The effect of GITC on RT-qPCR components was examined by amplifying beta-2-microglobulin (B2M) mRNA (13) in the dilution series of extracted RNA from OP and NP samples from a healthy individual, in an RT-qPCR run in two separate steps. The reverse transcription reaction (20 μL) contained 2 μL 10x M-MulV buffer (B0253S, New England Biolabs), 1 μL of 50 μM Oligo 18 dT (SO132, Thermo Fisher), 1 μL of 50 ng/μL random hexamers (18091050, Invitrogen), 1 μL of 10 mM of deoxynucleotide triphosphate (dNTPs) (180912050, Invitrogen), 0.2 μL of 40 U/μL RNase inhibitor (3335402001, Roche), 5 μL of the RNA extraction sample (diluted from 2x to 25x), 8.8 μL RNase-free H_2_O, and 1 μL of 200 U M-MuLV reverse transcriptase enzyme (B0253S, New England Biolabs). Negative controls were reactions without the RT enzyme and without sample, and the positive control was RNA isolated from HeLa cells. The RT-reaction was performed in Veriti™ 96-Well Thermal cycler (Applied Biosystems) (5 min at 25C, 1 h at 42C, and 20 min at 65C). The qPCR reaction (10 μL) contained 0.5 μL of the 10 μM B2M forward primer (5’-TGC CTG CCG TGT GAA CCA TGT-3’), 0.5 μL of the 10 μM B2M reverse primer (5’-TGC GGC ATC TTC AAA CCT CCA TGA-3’), 1 μL of the RT-reaction, 3 μL RNase-free H_2_O, 5 μL PowerUp SYBR Green Master Mix (A25741, Thermo Fisher). Reactions were set up in a 384-well plate and run in a QuantStudio 12K Flex Real-Time PCR System (4471081, Applied Biosystems) (2 min at 50C, 2 min at 95C, followed by 40 cycles of 95C for 15 sec; 60C for 1 min). Finally, a melting curve was recorded for 15 sec. at 95°C, 1 min at 60°C and 15 sec at 95°C. Data were analysed using QuantStudio™ 12K Flex Software. To confirm that only mRNA was amplified, the reactions were analysed by gel electrophoresis (data not shown). The B2M qPCR reaction proceeds with a forward primer placed in exon 2 and a reverse primer spanning the exon 3/4 junction to avoid amplification of the genomic B2M gene. With B2M cDNA, the primers produce an amplicon of 97 nucleotides, whereas the amplicon from the B2M gene itself spans 1974 nucleotides.

### One-step RT-qPCR against SARS-CoV-2

The effect of GITC on a corresponding one-step SARS-CoV-2 RT-qPCR reaction was examined using the dilution series of extracted RNA from OP and NP samples from the COVID-19 patient. SARS-CoV-2 specific RT-qPCR was also performed on samples from three previously SARS-CoV-2 positive, and a true negative, individual, to examine if non-specific amplification might occur. Three negative controls were used; NP and OP samples from a healthy individual and a test without sample. A primer and detection probe set against human RNase P mRNA was used as an internal positive control. The RT-qPCR was performed using SuperScript™ III One-Step RT-PCR System with Platinum™ Taq DNA Polymerase (12574-026, Invitrogen). Each 25 μL reaction contained 5 μL of sample, 12.5 μL of the 2x Superscript™ III reaction mix (a buffer containing 0.4 mM of each dNTP and 3.2 mM MgSO_4_), 0.4 μL 50 mM MgSO_4_, 0.05 μL of 20 μg/μL (1 μg BSA/reaction) of nonacetylated bovine serum albumin (BSA, 10711454001, Roche), 1 μL SuperScript™ III RT/Platinum *Taq* Mix and 1.85 μL of primer/probe mix (2019-nCoV CDC EUA Kit, 10006606, IDT). Primer and probe concentrations are outlined in **Table 1**. The thermal cycling conditions were: 55°C for 30 min, 95°C for 3 min, then 45 cycles of 95°C for 15 sec; 62°C for 30 sec; 68°C for 30 sec, followed by a final 68°C elongation step for 5 min on QuantStudio 12K Flex Real-Time PCR System (4471081, Applied Biosystems). Data were analysed using QuantStudio™ 12K Flex Software.

**Table 1.**
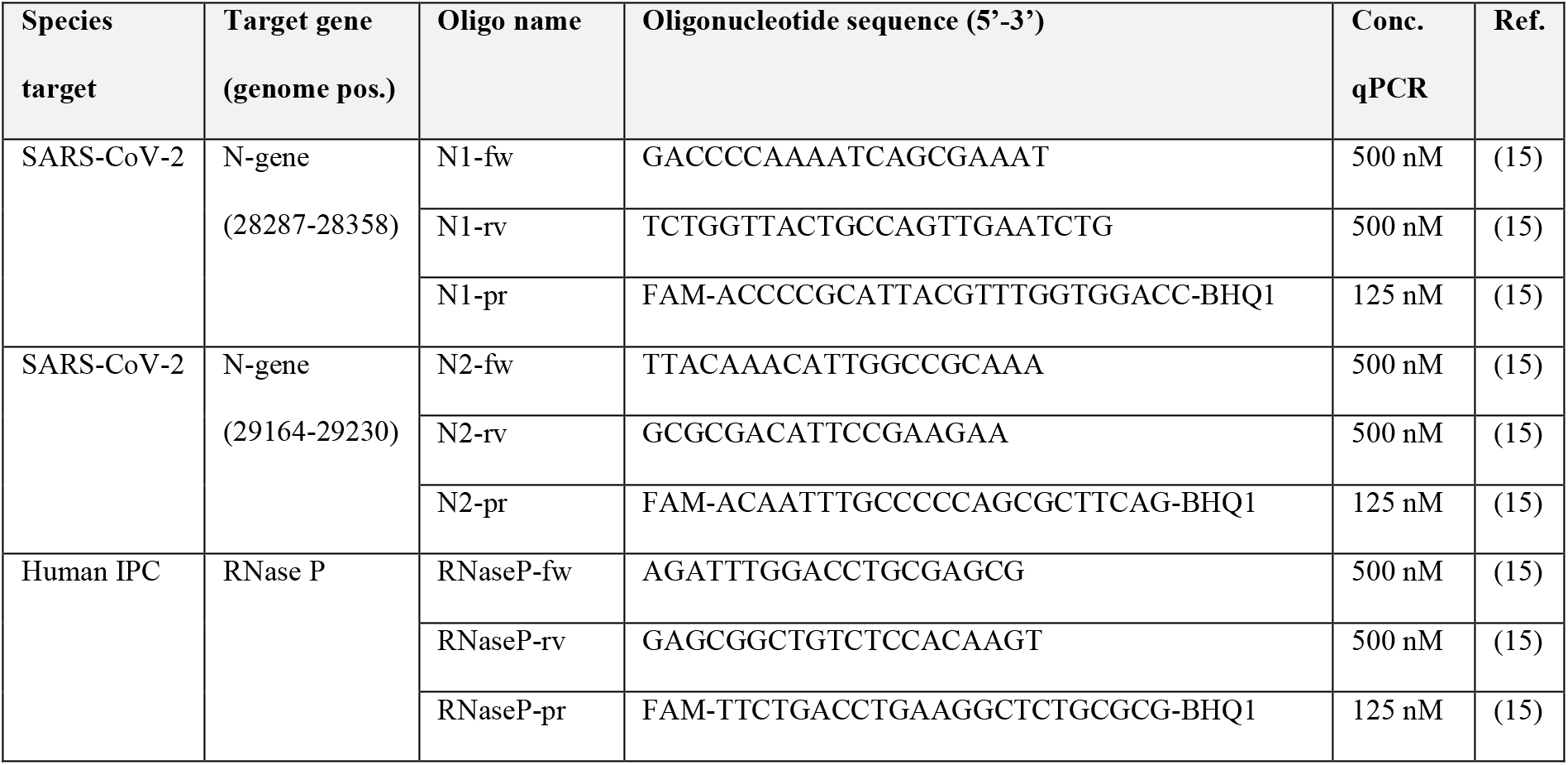
Overview of primers and probes used for SARS-CoV-2 detection. IPC: internal positive control, fw: forward primer, rv: reverse primer, pr: probe. Genome pos: genome position according to SARS-CoV-2 GenBank NC_004718.

### Preparation of a SARS-CoV-2 RNA dilution series using the GPC-extraction method

An OP sample swab was collected from a healthy individual and processed as described above. From the upper aqueous phase, 7.5 μL was carefully transferred to a new tube without disturbing the interphase, and spiked with 2.5 μL of Twist Synthetic SARS-CoV-2 RNA Control 1 (MT007544.1) to make a stock of 250.000 RNA copies/μL. This stock solution was used in a serial dilution with the remaining, un-spiked, aqueous phase to create a series of positive controls containing 3125, 781, 195, 48, 12, 3 or 0.76 RNA copies/μL. From each of these, 5 μL was used in a 25 μL one-step RT-qPCR reaction thus achieving a final number of 15625, 3906, 977, 244, 61, 15 and 3.8 SARS-CoV-2 RNA copies per one-step RT-qPCR reaction.

## RESULTS

### GITC/RNA dilution thresholds compatible with efficient two-step RT-qPCR

To investigate how GITC affected the efficiency of each step of an RT-qPCR reaction, we set up two-step B2M RT-qPCRs in the presence of variating concentrations of GITC. The B2M mRNA was extracted from OP and NP swabs from a healthy individual to mimic how patient material is obtained for SARS-CoV-2 testing. First, swabs were treated with the GPC solution. After mixing and centrifugation, this solution separated into an upper aqueous phase containing RNA, GITC and other salts, an interphase and a lower organic phase both containing DNA and proteins. The upper RNA/GITC aqueous phase was retrieved and diluted and used directly in an RT reaction at different final dilutions (ranging from 8x to 100x) followed by detection of B2M by qPCR with SYBR green (**Figure 1**). Consistent with GITC being a concentration-dependent mixed inhibitor (14), amplicons only appeared in the qPCR when the initial RT step was conducted with GITC dilutions ≥50x (**Figure 1**). Using the NP swab samples, amplicons were detected in the qPCR reaction with cycle threshold (Ct) values of 30-31 at a 50x GITC dilution. At the higher GITC dilution (100x), amplicons were detected at a later time with Ct-values > 35, likely due to the increased dilution of the B2M mRNA template in these reactions. When OP swabs were used, amplicons were similarly detected at ≥50x GITC dilutions (Fig. 1), but with a slightly higher Ct-values of 36. This difference in Ct thresholds for NP and OP swabs samples likely reflects differences in the amounts of cells obtained by the different sampling methods. Together, these results confirm that the RT-qPCR can proceed in the presence of low concentrations of GITC. The size of the B2M amplicons was examined using gel electrophoresis, and all amplicons were found to have the expected size of 97 nucleotides; no amplification of the genomic B2M gene was observed, and no amplicons were present in reactions without RT or sample. These results confirmed that the simplified GPC-extraction method is capable of providing RNA that can serve as a template in RT-qPCR.

### GITC/RNA dilution thresholds compatible with efficient one-step SARS-CoV-2 RT-qPCR

We next tested if the simplified GPC-extraction method was compatible with detection of SARS-CoV-2 in a hospitalised COVID-19 patient who had tested virus-positive three weeks earlier using the cobas^®^ SARS-CoV-2 test (Roche diagnostics). OP and NP samples were obtained from the patient, three former COVID-19 patients, and a healthy individual. For SARS-CoV-2 detection, we used a one-step RT-qPCR (11) with two different sets of primers and detection probes (N1 and N2) (15) (**Table 1**) specific for the conserved SARS-CoV-2 nucleoprotein (N) gene.

RNA was extracted from the OP and NP swabs and based on the B2M RT-qPCR results, the aqueous phase with RNA and GITC was used at a 50x, 75x, or 100x final dilution in the one-step SARS-CoV-2 RT-qPCRs. The OP swab sample from the COVID-19 patient yielded Ct-values of 27 at the 100x dilution for both duplicates for the N1 primer set, and Ct-values of 31 for both duplicates for the N2 primer set (**Figure 2a**). The RNase P internal control PCR resulted in Ct-values of 30 for both duplicates. No amplification was detected using the NP sample, which could reflect intermittent, low and/or no viral shedding from NP cells at this time (11), or problems with sampling from the nasopharynx. The three former COVID-19 patients who tested negative for SARS-CoV-2 using the cobas^®^ SARS-CoV-2 test on the day of comparison, and the healthy control, were virus-negative and RNase P positive. These results confirmed that the simplified GPC-extraction method, combined with a one-step RT-qPCR reaction, can be used to detect SARS-CoV-2 in an infected individual and that unspecific amplification does not seem to occur.

**Figure 2.**
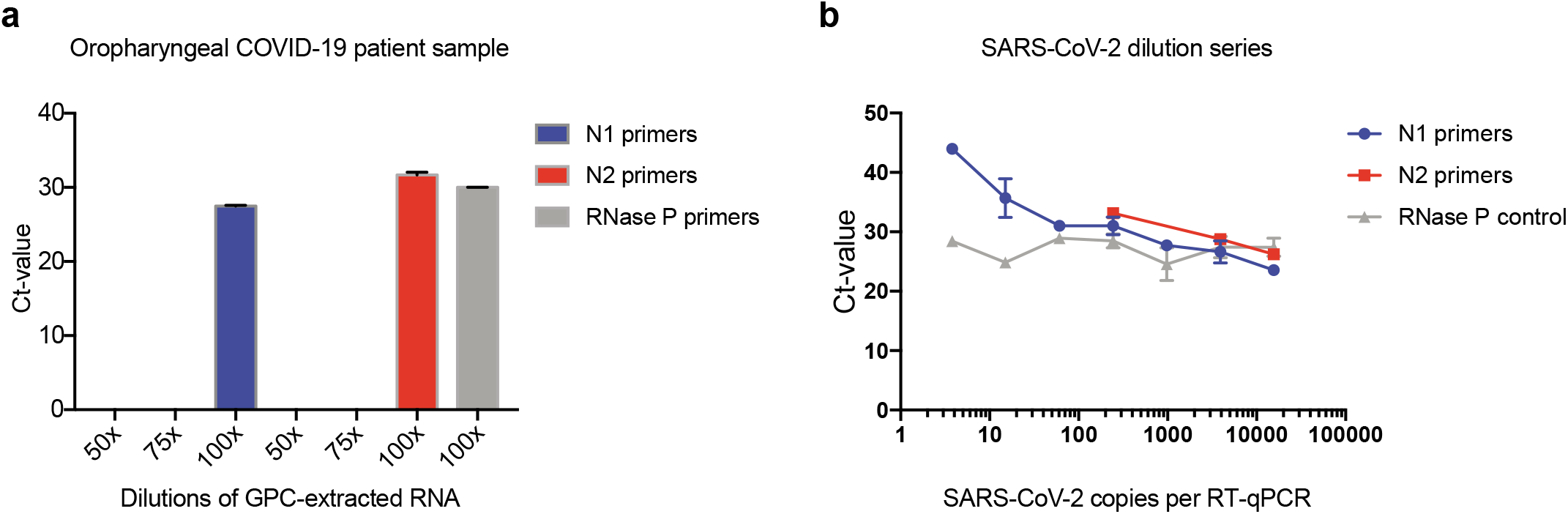
The effect of GITC and other salts on the efficiency of SARS-CoV-2 RT-qPCR and assay sensitivity. **a.** The effect of GITC and other salts on SARS-CoV-2 RT-qPCR detection of virus in the diluted aqueous phase of samples from a COVID-19 patient. Cycle threshold (Ct) for RT-qPCRs targeting the N-gene on the SARS-CoV-2 virus RNA, with viral RNA extracted from an OP swab from confirmed COVID-19 patient. The internal control RNase P RT-qPCR amplifications were negative at 50x and 75x dilutions (not shown). **b.** Determination of the sensitivity of the SARS-CoV-2 one-step RT-qPCR protocol using synthetic SARS-CoV-2 RNA combined with RNA extracted from an OP swab from a healthy individual using the GPC-extraction method. The concentration of GITC and other salts were constantly kept at a 100x dilution in each of the diluted virus samples. Cycle threshold (Ct) for RT-qPCRs targeting the N-gene on the synthetic SARS-CoV-2 virus RNA using the N1 and N2 primer sets, respectively. Amplification of RNAse P was used as an internal control. The amplifications were performed in duplicate.

### The simplified GPC-extraction allows detection of ~4 copies of SARS-CoV-2 per RT-qPCR

To assess the sensitivity of the COVID-19 diagnostic test flow, the one-step RT-qPCR was performed on a dilution series of a synthetic SARS-CoV-2 control RNA. RNA from an OP swab from a healthy individual was extracted using the simplified GPC-extraction method, after which some of the aqueous phase was spiked with the SARS-CoV-2 synthetic RNA to a final concentration of 250000 copies/μL. This spiked sample was used to create a dilution series ranging from 15625 to 3.8 copies/μL using the un-spiked aqueous phase from the sample as the diluent to retain consistent amounts of GITC and swab components in the RT-qPCRs. In accordance with the COVID-19 diagnostic test flow, each spiked sample was used at a final GITC dilution of 100x and SARS-CoV-2 RNA was detected with N1 or N2 primers, and using RNase P primers as positive control.

Amplicons were detected in all dilutions of the SARS-CoV-2 synthetic RNA down to 3.8 copies with the N1 primers (**Figure 2b**). This sensitivity is consistent with that reported for other primers that target the N gene (N-Sarbeco, Tib-Molbiol, Berlin, Germany) that showed a detection limit of 8.3 copies/reaction when used with a commercial RNA extraction kit (MagNA Pure 96 system, Roche, Penzberg, Germany) and the same one-step RT-qPCR (11). The ability to detect ~4 copies of viral RNA in the RT-qPCR reaction translates into ~10^4^ viral copies per swab, which is more than 10 fold lower than the average virus load per NP or OP swab from symptoms onset to day five (6.76×10^5^ copies per swab) and later (3.44×10^5^ copies per swab) (10, 11). The N2 primers proved less sensitive, detecting synthetic RNA down to only 244 copies/reaction in our set-up. It cannot, however, be excluded that the sensitivity is nearer the next testing point of 61 copies/reaction. The negative control with no added virus showed no amplification, whereas efficient amplification was observed with all positive control reactions targeting RNase P. Together, these data demonstrate that the simplified GPC-extraction method allows for similar detection sensitivity in the one-step RT-qPCR as a currently utilised kit-based RNA extraction methods.

Based on combined experiments, we outlined a COVID-19 diagnostic test flow from patient-to-result that covers patient sampling, RNA extraction by the simplified GPC-extraction method, and one-step RT-qPCR detection (**Figure 3**).

**Figure 3.**
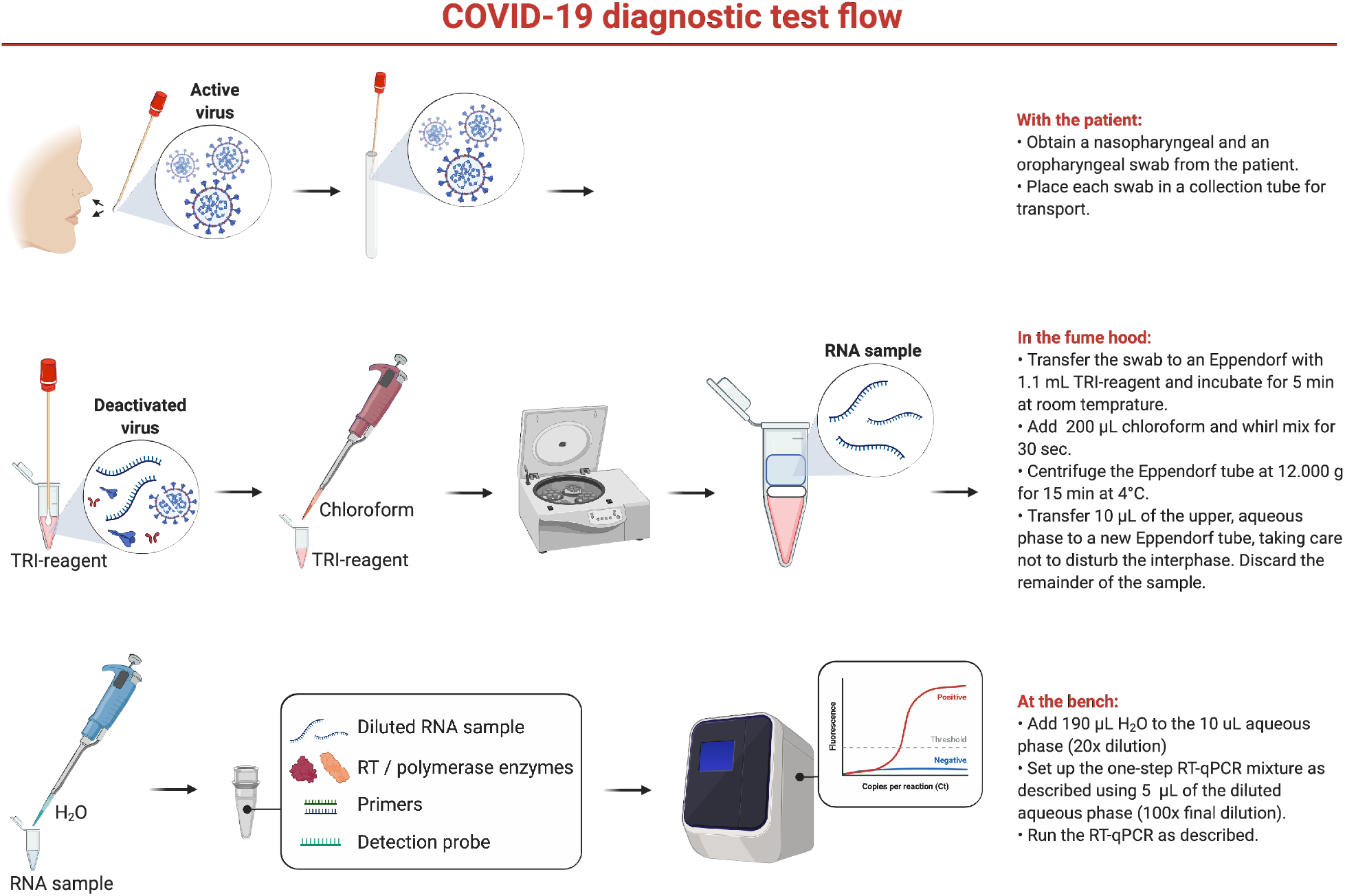
Workflow for SARS-CoV-2 detection using the simplified GPC-extraction method.

## DISCUSSION

The ability to test large segments of the population represents the most effective means of managing the SARS-CoV-2 pandemic and making informed decisions on the reopening of our societies. To date, such tests typically combine the use of a front-end RNA extraction kit and a one-step RT-qPCR detection kit. Many commercial RNA extraction kits have been shown to not fully inactivate the virus, potentially putting healthcare personnel that handles the samples at risk of SARS-CoV-2 infection (18). Similarly, several recently published quick RNA-extraction methods (7, 16, 17) that rely on inactivation of the virus by heat, do not completely inactivate the virus (18). In contrast, the GPC-solution fully inactivates corona viruses such as MERS-CoV (12), adding a desirable safety aspect to this method.

The GITC salt is a mixed inhibitor of PCRs, affecting both the function of polymerase enzymes and the melting temperature of primer/template duplexes (14). GITC is therefore usually removed from the RNA by precipitation of the RNA with isopropanol, centrifugation and washing of the RNA pellet with 70% ethanol after which the pellet is dissolved in H2O. This process requires expertise in RNA handling as GITC can co-precipitate with the RNA, and too vigorous pipetting or wrongly handled RNA pellet solvation can result in RNA loss. A previous study demonstrated that GITC had only a modest inhibitory effect at a concentration of 9 μg/μL (~75 mM) (14), prompting us to speculate that the RNA/GITC aqueous phase could be used directly in RT-qPCR without precipitation, centrifugation, washing and solvation if the solution was diluted below the ~75mM threshold. Our results robustly demonstrate that a dilution of 50x and 100x of the aqueous phase is compatible with one- and two-step RT-qPCRs, respectively, and that approximately ~4 copies of SARS-CoV-2 can be detected, equivalent to ~10^4^ virus copies per NP or OP swab. The difference between the one- and two-step RT-qPCRs results are likely due to the extended period of exposure of the PCR polymerase to GITC salts during the latter procedure.

As the simplified GPC-extraction method presented here rapidly inactivates the virus, it allows detection by RT-qPCR in laboratories not classified to handle infectious air-borne viruses. The GCP-solution also denatures proteins to prevent enzymatic degradation of the viral RNA genome, which facilitates sample storage prior to extraction when needed. Moreover, the simplified GCP-extraction method utilises equipment and reagents common to clinical and molecular biology laboratories, thus removing the reliance on commercial RNA-extraction kits that presents a bottleneck for large-scale SARS-CoV-2 testing. The simplified GPC-extraction method shortens the time from patient sampling to testing relative to using the full GPC-extraction protocol (19), but, more importantly, excludes the steps that require experience with RNA precipitation, washing and reconstitution. The resulting lowered complexity makes this simplified method amenable to non-RNA-experts, thereby increasing the number of laboratories at which these tests can be performed. The sensitivity of the downstream RT-qPCR was similar to that reported previously (11) and appears superior to that of other RT-qPCR protocols, including the cobas^®^ SARS-CoV-2 test (20). The difference in sensitivity between the N1 and N2 primer sets is important when evaluating the result of SARS-CoV-2 test using both primers sets as confirmatory for the infection, since a negative result for the N2 primers, but positive result for the N1 primers, could simply reflect a lower viral load in the tested infected individual, compared to a patient with two positive results.

In the here-presented protocol we add the virus-inactivating TRI-reagent after the collected samples are transferred to the laboratory, where this solution can be safely handled in a fume hood. It is tempting to speculate that a test tube could be developed that enables contact between the swab and the TRI-reagent at the point of sampling, without exposing testing personal to the reagent, thus inactivating the virus at the earliest possible time point in the diagnostic procedure.

In summary, our protocol for RNA extraction relieves the dependence on expensive commercial kits that have become a bottleneck in the diagnosis of the virus, and ensures the safety of healthcare workers testing for SARS-CoV-2 infections, which in turn expands the number of testing laboratories and thus testing capacity.

## ACKNOWLEDGEMENT

We are grateful to Camilla Thomsen for sample handling and processing.

## FUNDING STATEMENT

No external funding was obtained for this work.

## AUTHOR DECLARATIONS

The authors declare no competing interests.

Standard ethical guidelines and regulations regarding method development and optimisation were followed. All necessary patient/participant consent has been obtained and the appropriate institutional forms have been archived. No project-specific ethical approval was necessary.

All data are available from the authors upon request.

